# Spatial Transcriptomics in Bone Mechanomics: Exploring the Mechanoregulation of Fracture Healing in the Era of Spatial Omics

**DOI:** 10.1101/2024.04.18.590091

**Authors:** Neashan Mathavan, Amit Singh, Francisco Correia Marques, Denise Günther, Gisela Kuhn, Esther Wehrle, Ralph Müller

## Abstract

In recent decades, the field of bone mechanobiology has sought experimental techniques to unravel the molecular mechanisms governing the phenomenon of mechanically-regulated fracture healing. Each cell within a fracture site resides within different local micro-environments characterized by different levels of mechanical strain - thus, preserving the spatial location of each cell is critical in relating cellular responses to mechanical stimuli. Our spatial transcriptomics based “mechanomics” platform facilitates spatially-resolved analysis of the molecular profiles of cells with respect to their local *in vivo* mechanical environment by integrating time-lapsed *in vivo* micro-computed tomography, spatial transcriptomics, and micro-finite element analysis. We investigate the transcriptomic responses of cells as a function of the local strain magnitude by identifying the differential expression of genes in regions of high and low strain within a fracture site. Our platform thus has the potential to address fundamental open questions within the field and to discover mechano-responsive targets to enhance fracture healing.

## Introduction

In the late 19th century, the German orthopedic surgeon Julius Wolff established the fundamental principle of bone mechanobiology by describing the dynamic nature of bones and their remarkable ability to adapt to their mechanical environment (*1*). This principle underscores the critical importance of the mechanical environment to the fracture healing capacity of bone. Mechanical stimuli can either enhance or impair the fracture healing process (*2*). At its core, the mechanobiology of fracture healing is governed by the response of cells at the fracture site to physical stimuli. However, it has proven extremely difficult to investigate the transduction of mechanical stimuli exerted at the organ level to site-specific cellular responses at the molecular level. Despite considerable advances in recent decades (*3*, *4*), our understanding of the specific signaling pathways underlying the mechanoregulation of fracture healing remains in its nascent stages.

Each cell within a fracture site resides within different local micro-environments characterized by different levels of mechanical strain. The challenge within the field has been the need for more integrative approaches to investigate the multi-scale repair response of individual cells in response to their local *in vivo* mechanical environment (*5*). Insights into this critical missing link have the potential to be transformative within the field by greatly improving our ability to anticipate cellular responses to different magnitudes or modes of mechanical stimuli. In pursuing the characterization of cellular activity as a function of its local *in vivo* mechanical environment, experimental approaches have largely been confined to *in vitro* (*6*) and *in silico* (*5*) techniques. The challenges with *in vivo* approaches are substantial. Rodent fracture models (*2*), osteogenic loading protocols (*7*) and *in vivo* imaging techniques (time-lapsed micro-computed tomography, micro-CT) (*8*) are well-established within the field. However, *in vivo* mechanical loading applied at the organ scale is heterogeneously distributed throughout the fracture site, resulting in complex mechanical environments with distinct regions of high and low strain. Spatial correlation is thus required to associate cellular responses with their respective local *in vivo* mechanical environments. Currently, micro-finite element analysis (micro-FE) is the only established technique available to generate 3D maps of the *in vivo* mechanical environment. High-resolution micro-FE models derived from micro-CT images of the fracture site have been used to associate morphological changes at the tissue scale with the local mechanical environment (*9*). Moreover, osteocytes – the primary mechanosensory cell in bone and constituting > 90% of all bone cells – reside deep within the bone matrix. Direct experimental observation of these cells is thus challenging without destruction of or interference with the surrounding tissue environment. Mechanomics – the application of omics technologies to investigate the interactions between local mechanical environments and cellular/molecular responses – holds immense potential but is technically challenging. Techniques such as laser capture microdissection, in combination with FE modelling, have permitted “mechanomic” analyses of a small number of isolated cells (*5*). In contrast, recent advances in spatially resolved “omics” technologies now permit the comprehensive, unbiased mapping of molecular pathways and cellular function within the spatial context of complex tissue architectures (*10*). However, the use of spatial technologies in bone has been limited due to the calcified nature of the tissue.

To investigate the molecular responses of cells to their local mechanical environment within a mechanically loaded fracture site, we have established a spatial-transcriptomics based “mechanomics” platform (fig 1). Spatial transcriptomics – a recent, transformative advancement in cellular profiling technologies – permits characterization of the transcriptomic responses of cells within their native spatial context on histology sections. Our platform consists of: (i) an established femur defect mouse model (*11*), (ii) established *in vivo* micro-CT imaging protocols and analyses (*9*), (iii) an established osteogenic cyclic mechanical loading protocol (*11*, *12*), (iv) an established spatial transcriptomics protocol for bone tissue (*13*) and (v) an established *in silico* micro-FE modelling approach (*12*). As each component of our platform has been previously established, these techniques, models and protocols collectively provide a solid scientific foundation for our platform. Our objective is to present a "proof-of-principle" study to demonstrate the potential of a spatial transcriptomics-based “mechanomics” platform to identify the molecular mechanisms governing the mechanoregulation of fracture healing. As illustrated in Figure 1, we introduced femoral osteotomies in mice and performed weekly *in vivo* micro-CT imaging of the fracture sites. Following bridging of the fracture site at 3 weeks post-surgery, the mice were subdivided into Control and Loaded groups and received either sham loading or cyclic mechanical loading. Our analysis of the response to loading was based on Wolff’s Law – the cornerstone of the field of bone mechanobiology – which asserts that bone adapts to its mechanical environment by forming bone at sites of mechanical loading and resorbing bone at sites of mechanical unloading. Bone formation and resorption responses at the fracture site were thus the focus of our analyses. We first quantified the effects of mechanical loading using micro-CT-derived bone morphometric indices and visualized the sites of bone formation, quiescence and resorption. Next, we corroborated our findings by comparing the transcriptomic responses in Control vs. Loaded fracture sites. To correlate spatially resolved gene expression profiles with their local *in vivo* mechanical environments, we analyzed the transcriptomic profiles at sites of high strain and low strain within a mechanically loaded fracture site. Finally, we assessed the merits of our spatial mechanomics platform and its potential to develop a molecular-based understanding of the local mechanoregulation of fracture healing.

**Fig. 1.**
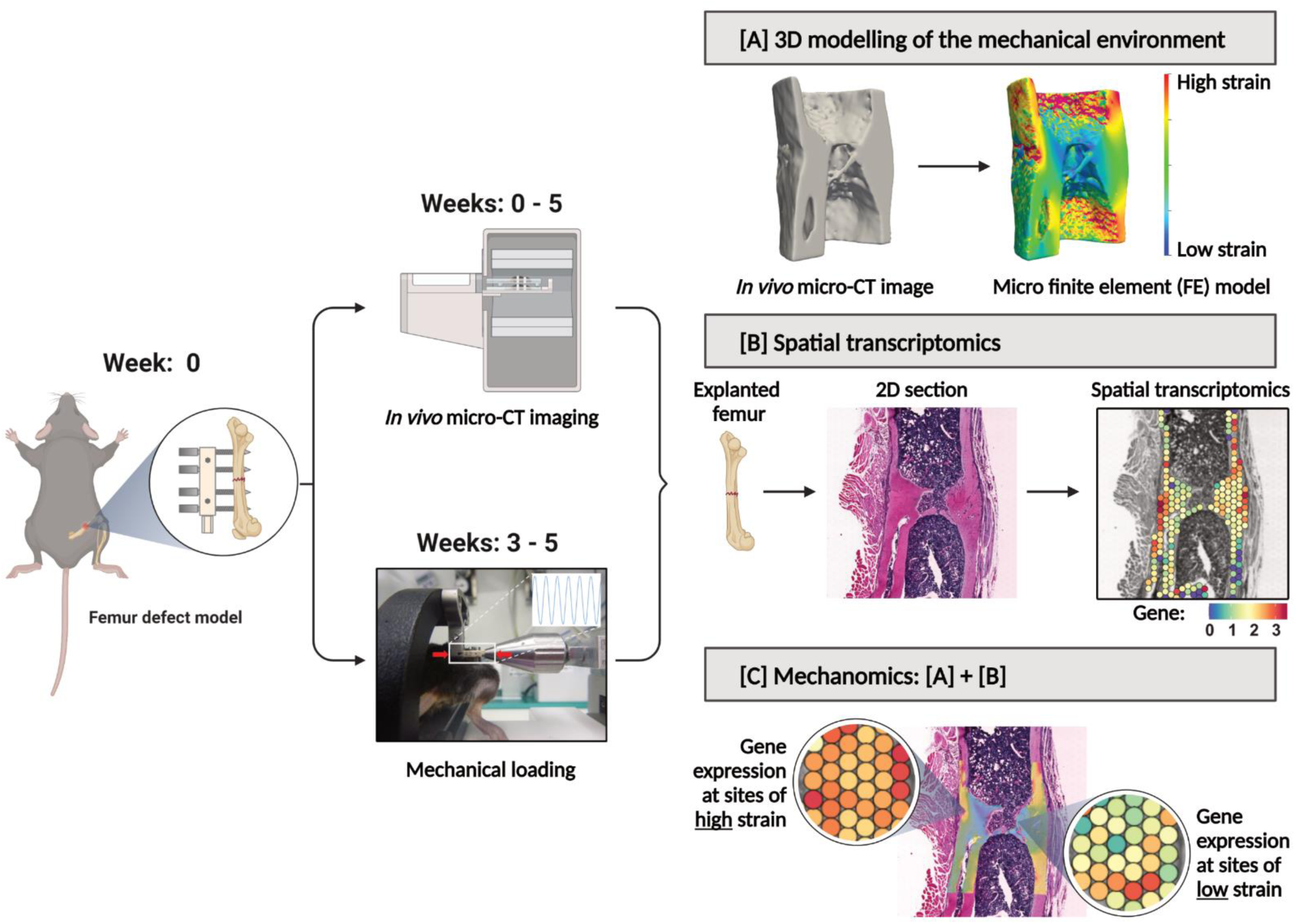
Overview of our spatial transcriptomics-based “mechanomics” platform to investigate the mechanobiology of fracture healing. The platform permits the generation and spatial integration of multi-modal data sets (CT bone morphology data, 3D mechanical environments and spatially resolved gene expression data) from a single fracture site. At week 0, mid-diaphyseal femoral osteotomies are introduced in the right femur of mice and stabilized with an external fixator. Time-lapsed *in vivo* micro-CT imaging is performed weekly at the fracture site (weeks 0- 5; 10.5 μm resolution). Mice which exhibit bridging at 3 weeks post-surgery are subdivided into Loaded and Control groups. At weeks 3 – 5, mice received individualized cyclic loading (up to 16N) or 0N sham-loading three times per week. All mice are euthanized at 5 weeks post-surgery. **[A]** Micro-FE analyses based upon *in vivo* micro-CT images are used to generate tissue-scale 3D maps of the mechanical environment. **[B]** Spatial transcriptomics analyses are performed on explanted femurs. To associate spatially resolved molecular profiles of cells with their local *in vivo* mechanical environment, the spatial transcriptomics histology section is visually aligned within the 3D map of their mechanical environment. **[C]** Gene expression can thus be analyzed as a function of the local mechanical environment. Illustration created with BioRender.

## Results

### Micro-CT-based bone morphometric analysis underscores a strong anabolic response to cyclic mechanical loading

Time-lapsed *in vivo* imaging permits visualization of the sites of bone formation, quiescence and resorption at the fracture site as shown at weekly intervals in Figure 2. Comparable healing responses were observed between Control and Loaded fracture sites between weeks 0 – 3. This was reinforced by comparing bone morphometric parameters between weeks 0 – 3 across all volumes of interest (fig 3 [A] – [H]). Upon bridging at week 3, the fracture sites of Loaded mice were subjected to cyclic mechanical loading 3 times per week. Loading induced a strong anabolic response as can be observed in Figure 2 where sites of bone formation (in orange) predominate at the loaded fracture site between weeks 3 - 5. In contrast, the fracture sites of Control mice were observed to undergo remodeling during the same period (fig 2). In comparisons of bone morphometric parameters, cyclic mechanical loading was found to induce larger callus / bone volume formation. At week 5, BV/TV in the defect center was 41.5% and 40.4% in Control mice vs. 75.4% and 66.3% in Loaded mice (fig 3[A]). Similarly, in the defect periphery, BV/TV was 11.4% and 16.7% in Control mice vs. 38.6% and 25.3% in Loaded mice (fig 3[B]). At week 5, loading induced an increased rate of bone formation (0.58% and 0.88% per day in Control mice vs. 1.43% and 1.90% per day in Loaded mice) and a diminished rate of bone resorption (-0.84% and - 0.58% per day in Control mice vs. -0.11% and -0.11% per day in Loaded mice) in the defect center (fig 3[E]). Similarly, in the defect periphery, loading induced an increased rate of bone formation (0.25% and 0.18% per day in Control mice vs. 1.74% and 1.47% per day in Loaded mice) and a diminished rate of bone resorption (-0.21% and -0.70% per day in Control mice vs. -0.06% and - 0.03% per day in Loaded mice) (fig 3[F]).

**Fig. 2.**
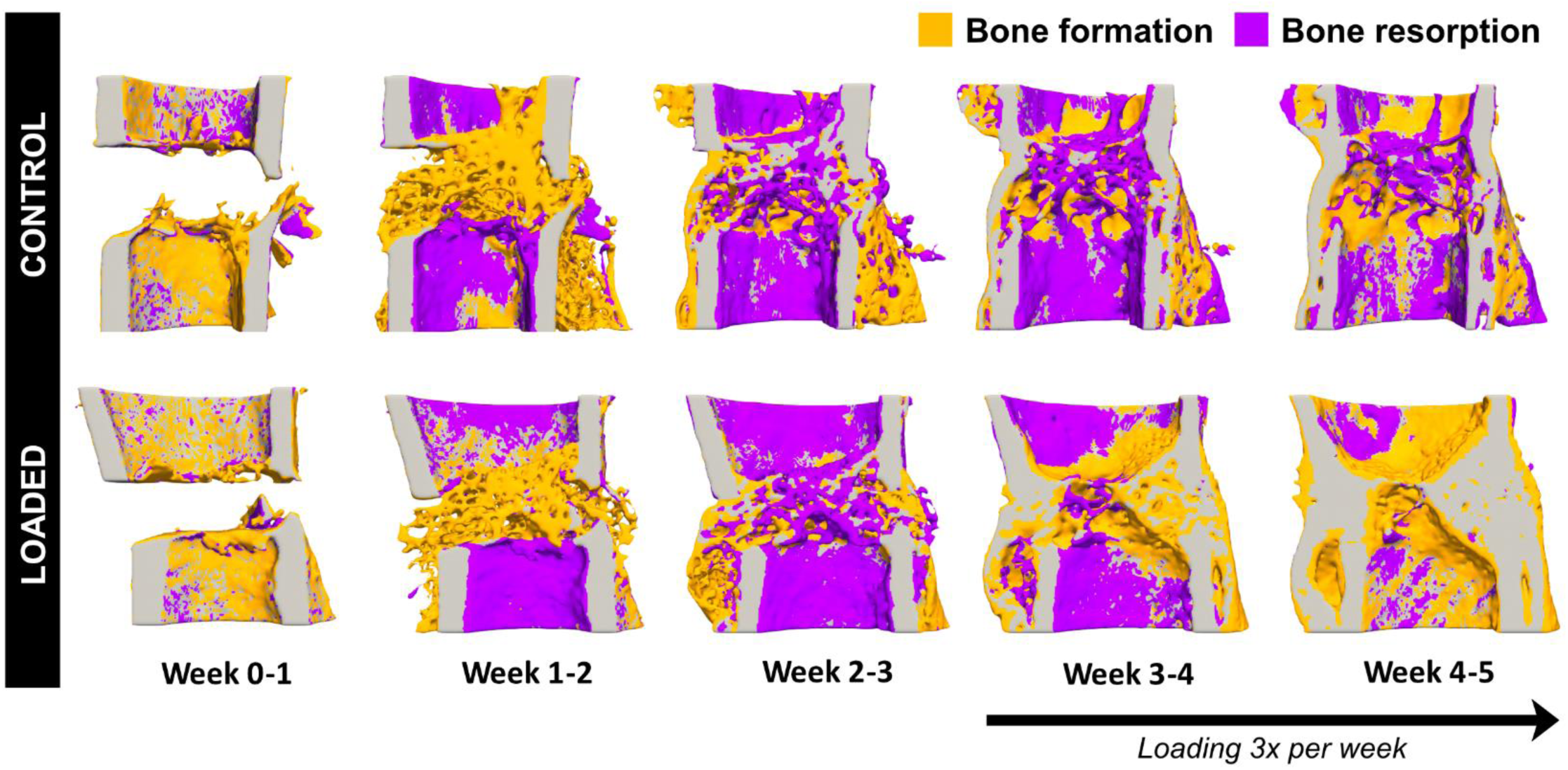
Visualization of sites of bone formation, quiescence and resorption in Control and Loaded fracture sites. Sites of bone formation (orange) and bone resorption (purple) are identified via registration of time-lapsed *in vivo* images (threshold: 395 mg HA/cm^3^, voxel size = 10.5 μm). In Loaded fracture sites, loading was applied 3x per week from Week 3 onwards. Visualization performed using Paraview (version 5.7.0). Spatial transcriptomics data generated from 2D sections of these samples are presented in Figure 4.

**Fig. 3.**
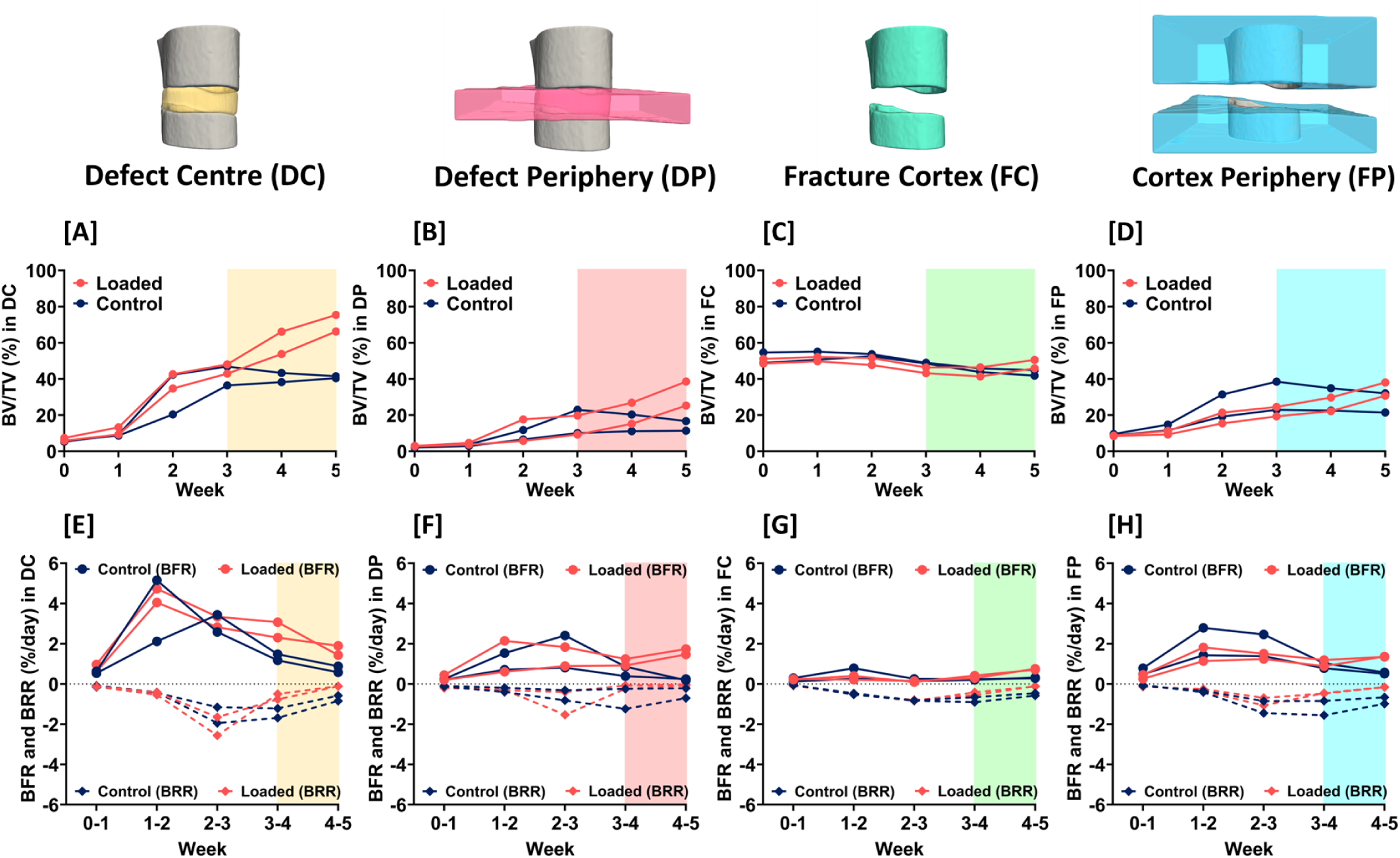
*In vivo* micro-CT morphometric analysis. Quantitative morphometric analyses (n = 2 per group) were performed in four volumes of interest: the defect center (DC), the defect periphery (DP), the existing fracture cortex and medullary cavity (FC) and the cortex periphery (FP). Two parameters are presented: **[A] – [D]**: BV/TV where the bone volume (BV) is normalized to TV (DC for DC and DP, FC for FC and FP). **[E] – [H]**: bone formation rates (BFR) and bone resorption rates (BRR). Shaded regions in each plot correspond to timepoints at which loading was applied 3x per week.

### Significantly enhanced osteogenic response corroborated by spatial analyses of differential gene expression

Spatial analyses of differential gene expression (DEG) in bone regions at the fracture sites of Control and Loaded mice revealed significantly higher expression of osteogenic markers in response to mechanical loading (fig 4-5). Regions for the analysis were defined by selecting spots in each histological section encompassing all bone spots at the fracture site between the two inner pins of the external fixator (fig 5[A]). Quality control measures for these bone regions were as follows: the median number of unique molecular identifiers (UMIs) for the Control and Loaded sections were 3602 and 2708 respectively, and the median number of unique genes for the Control and Loaded sections were 2670 and 1924 respectively. In these defined bone regions of the fracture site, 9889 genes were identified with sufficiently high expression to be included in the DEG analysis. Of these, 834 genes were differentially expressed (FDR-adjusted p value cutoff < 0.05; absolute log2 fold change > 0.5) with 395 genes upregulated and 439 genes downregulated (table S2). Furthermore, we ranked the genes by log2 fold change and compared two FDR-adjusted p value thresholds (0.05 and 0.01) as a measure of the consistency of the most regulated genes (table S2). We found the top 11 genes were identical at both thresholds – indicating that these genes are strongly associated with the response to mechanical loading. To present differential gene expression in subsequent sections, the following convention is used: *Gene* (+/- log2 fold change).

**Fig. 4.**
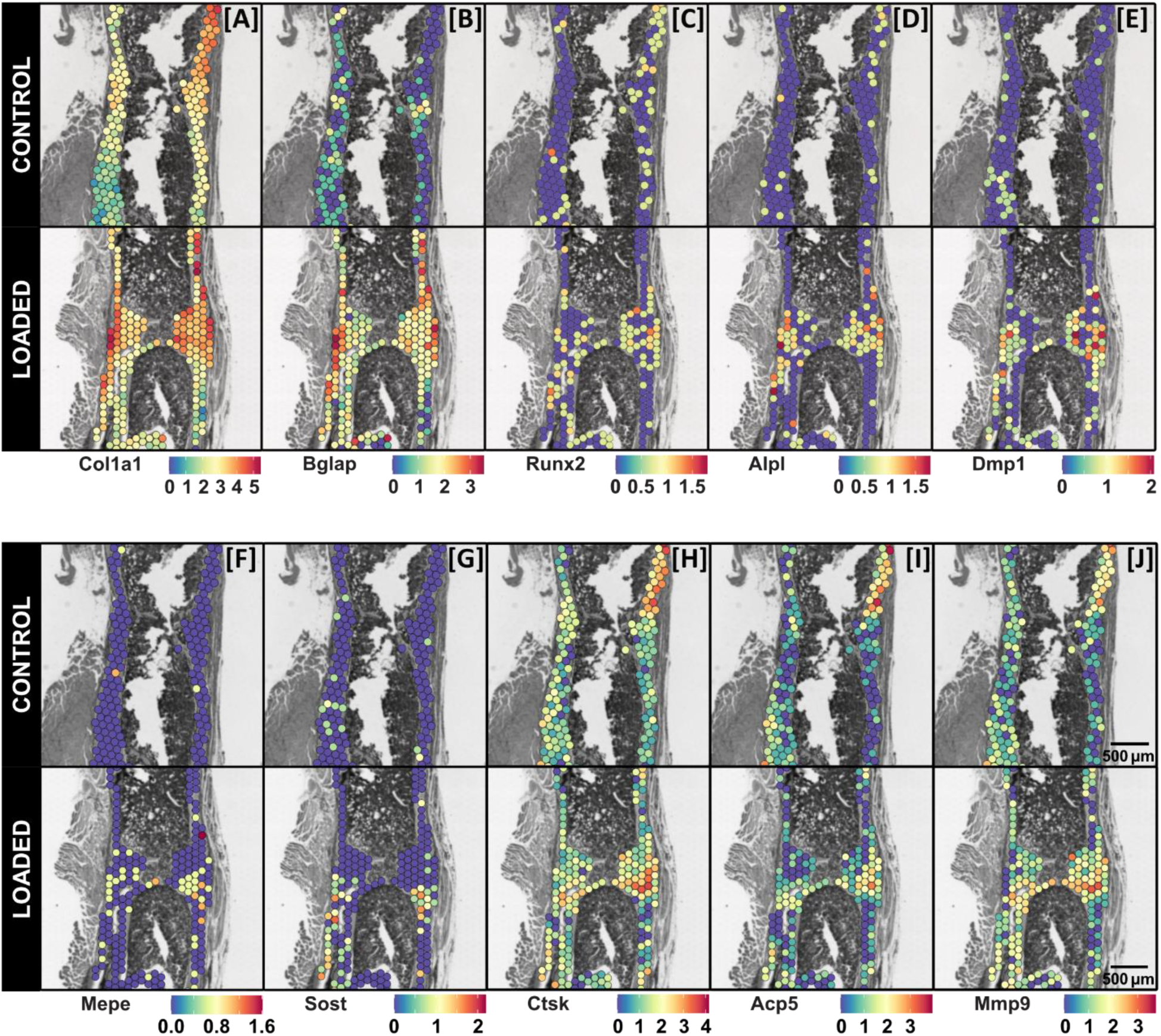
Spatial gene expression maps of selected bone cell markers at the fracture site of Control and Loaded mice. Visualization of the spatial expression patterns of osteoblast markers: [A] *Col1a1*, [B] *Bglap*, [C] *Runx2*, [D] *Alpl*, osteocyte markers: [E] *Dmp1*, [F] *Mepe*, [G] *Sost*, and osteoclast markers: [H] *Ctsk*, [I] *Acp5*, [J] *Mmp9*, within the fracture sites of Control and Loaded mice are presented. Each legend denotes the normalized expression of the specified gene. Data presented (n = 1 per group) corresponds to samples at 5 weeks post-surgery. 3D visualizations of the morphology of these Control and Loaded fracture sites are presented in Figure 2. Spatial transcriptomics spot size = 55 μm.

**Fig. 5.**
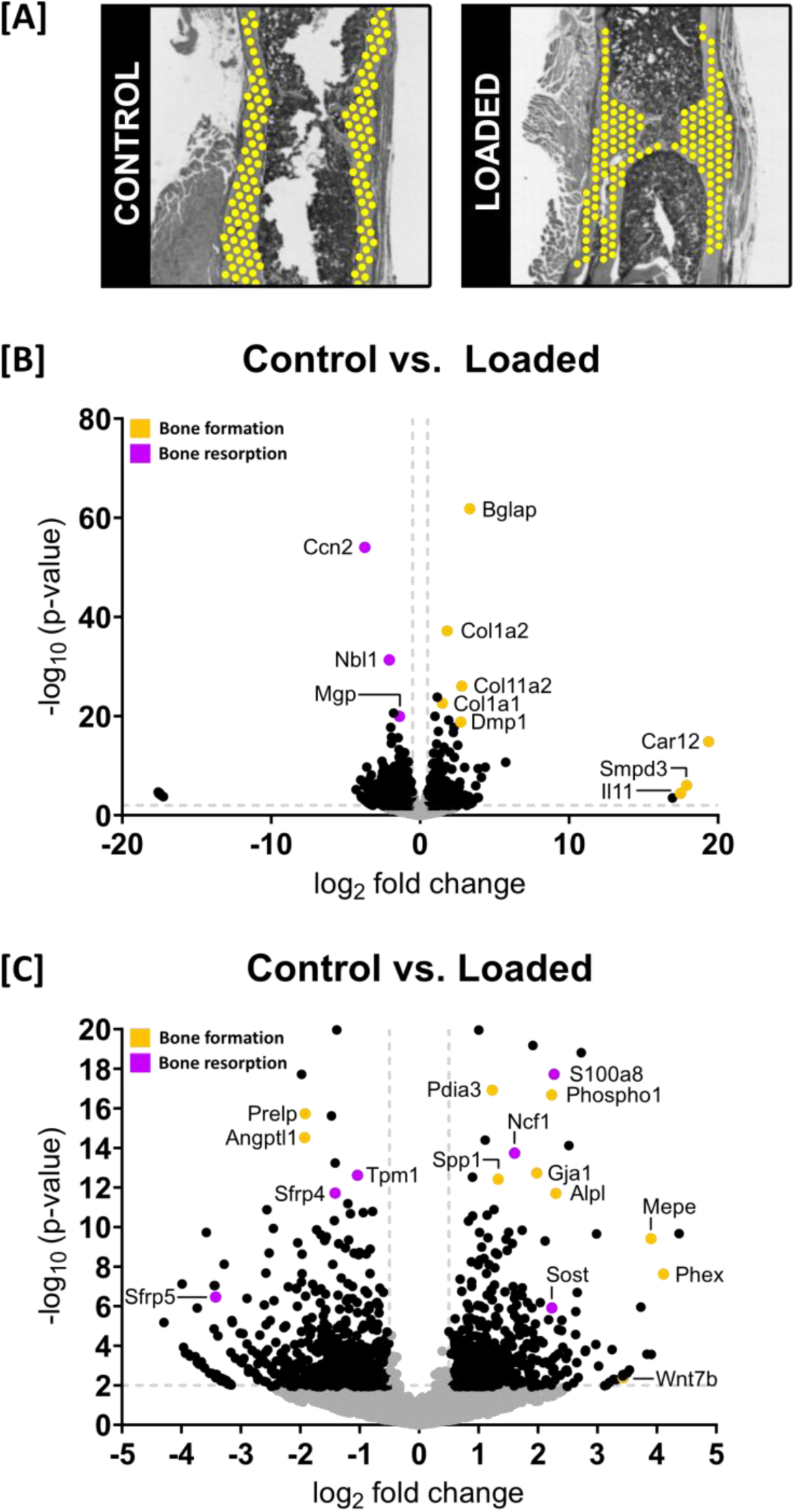
Gene expression profiling in Control vs. Loaded fracture sites. Data presented (n = 1 per group) corresponds to samples at 5 weeks post-surgery. **[A]**: Areas of interest defined in each fracture site for the analysis. **[B]**: Volcano plot to visualize differentially expressed genes (DEGs) (significance criteria: FDR-adjusted p value cutoff < 0.05 and an absolute log2 fold change > 0.5). Significant DEGs associated with bone formation are identified in orange and significant DEGs associated with bone resorption (or are inhibitors / antagonists of bone formation) are identified in purple. Non-DEGs are represented in gray. **[C]**: Magnified view of volcano plot to highlight DEGs of interest.

Significantly higher expression of markers of osteoblast differentiation and osteoblast activity were present in response to loading: *Bglap* (+3.3 fold), *Alpl* (+2.3 fold), *Sp7* (+1.7 fold), *Col1a1* (+1.5 fold), *Col1a2* (+1.5 fold), and *Runx2* (+0.9 fold) (fig 5[B][C]; table S2). Visualization of the spatial expression of these genes further underscored this result (fig 4 [A] – [D]). Markers of mineralizing osteocytes: *Phex* (+4.1 fold), *Dmp1* (+2.7 fold), and mature osteocytes: *Mepe* (+3.9 fold), *Sost* (+2.2 fold) were also upregulated (fig 4 [E] – [G]; fig 5[B][C]). Furthermore, amongst the top-ranked upregulated genes include genes associated with osteogenesis (*14*): *Wnt7b* (+3.4 fold), endochondral ossification (*15*, *16*): *Adam12* (+3.7 fold), matrix synthesis (*17*): *Cdo1* (+4.4 fold), and mineralization (*18*–*20*): *Car12* (+19.4 fold), *Smpd3* (+17.9 fold) and *Phospho1* (+2.2 fold) (fig 5[B][C]; table S2).

Coincident with this enhanced osteogenic response, *Ccn2* (-3.7 fold) – a master regulator of osteogenesis and chondrogenesis (*21*), *Nbl1* (-2.1 fold) – a BMP-antagonist (*22*), and *Mgp* (-1.4 fold) – a potent mineralization inhibitor (*23*, *24*), were downregulated (fig 5[B]; table S2). Osteoclast markers were less prominent in the results with expression of *Ctsk* and *Acp5* not significantly different between Control and Loaded fracture sites and upregulation of genes associated with osteoclastogenesis (*25*, *26*) and osteoclast activity (*26*–*29*): *s100a8* (+2.3 fold), *Ncf1* (+1.6 fold) and *Mmp9* (+0.8 fold) (fig 4 [H] – [J]; fig 5[B][C]; table S2). Cartilage was minimally present at both fracture sites, as evident in the gene expression maps for selected chondrocyte markers (*30*, *31*) shown in Supplementary Figure S1.

Known mechano-responsive genes were also present amongst the top-ranked genes including: *Il11* (+17.5 fold) – a cytokine predominantly expressed in bone (*32*, *33*), *Ccn2* (-3.7 fold) (*21*, *34*), *Wnt7b* (+3.4 fold) (*35*, *36*), *Sost* (+2.2 fold) – a Wnt-antagonist (*37*, *38*), *Gja1* (+2.0 fold) – which encodes for the gap junction protein connexin 43 (*39*), *Sfrp4* (-1.4 fold) – a Wnt-antagonist (*40*– *42*), and *Cdkn1a* (-0.7 fold) – a negative regulator of osteogenesis (*43*, *44*) (fig 5[B][C]; table S2).

Differential gene expression analyses between Control vs. Loaded sites were repeated with the region of interest narrowed to sites of newly formed bone (fig S2). Using the DC and DP volumes illustrated in Figure 3, spots were chosen which correspond to sites of newly formed bone (fig S2[A]). Differential gene expression was found to be largely similar to the results reported using the larger regions of interest (fig S2[B][C]).

### Mechanomics analyses reveal regions of high / low strain are associated with sites where bone formation / resorption responses respectively predominate

In the Loaded spatial transcriptomics section, gene expression profiles corresponding to high strain (effective strain or EFF > 1000 µε, n = 22 spots), low strain (EFF < 500 µε, n = 24 spots) and reference strain regions (EFF > 500 µε and EFF < 1000 µε, n = 42 spots) were analyzed (fig 7). Using the co-efficient of variance (CV) as a measure of functional significance, the top-ranked genes in regions of high strain included genes associated with an anabolic response: *Spp1*, *Col1a2*, *Col1a1*, *Bglap*, *Sparc*, *Col11a2*, *Gpx3*, *Pdia3* (*45*), the osteoclastogenesis inhibitor (*46*): *Cd74* and the mineralization inhibitor: *Mgp* (fig 8). In contrast, the top-ranked genes in regions of low strain included genes associated with a catabolic response: *S100a8*, *Mmp9*, *Tpm1* (*47*, *48*), *Ctsk*, *Ncf1* and *Igfbp4* (*49*, *50*) (fig 8). In the reference strain region, the top-ranked genes included genes associated with an anabolic response: *Spp1*, *Sparc*, *Bglap*; genes associated with a catabolic response: *Ctsk*, *Acp5*, *Mmp9*; and genes which couple bone formation with resorption: *Mmp13* (*27*) (fig 8).

**Fig. 6.**
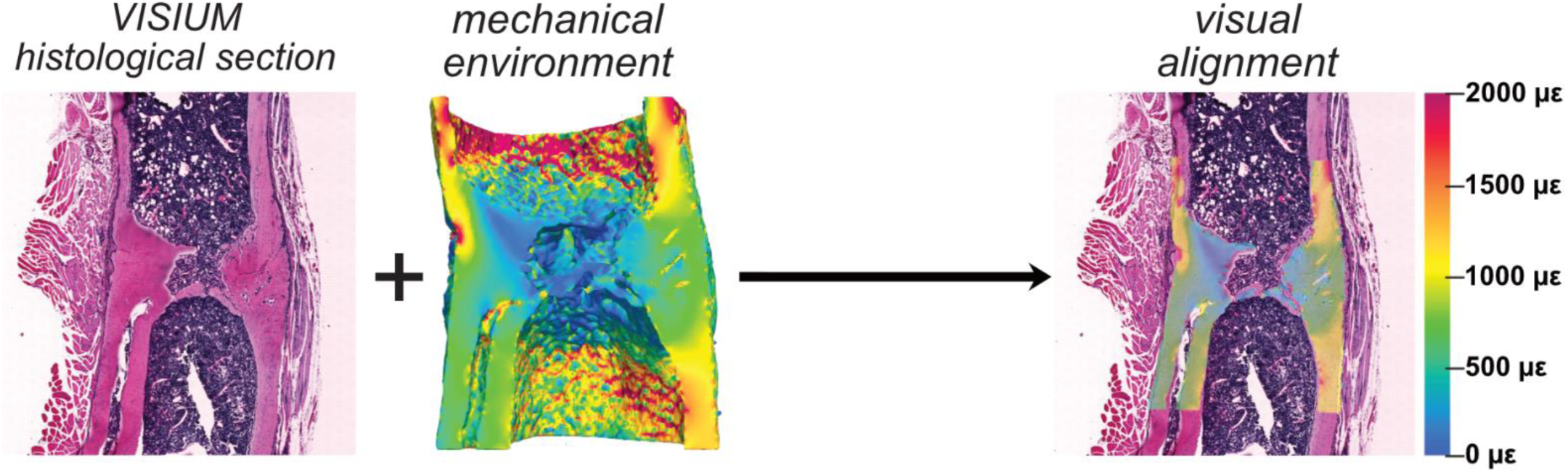
Association of spatially resolved molecular profiles of cells with their local *in vivo* mechanical environment. Visual alignment of the 2D spatial transcriptomics histological section within the 3D mechanical environment in the Loaded fracture site. Element size in the 3D micro-FE model of the mechanical environment is 10.5 x 10.5 x 10.5 μm.

**Fig. 7.**
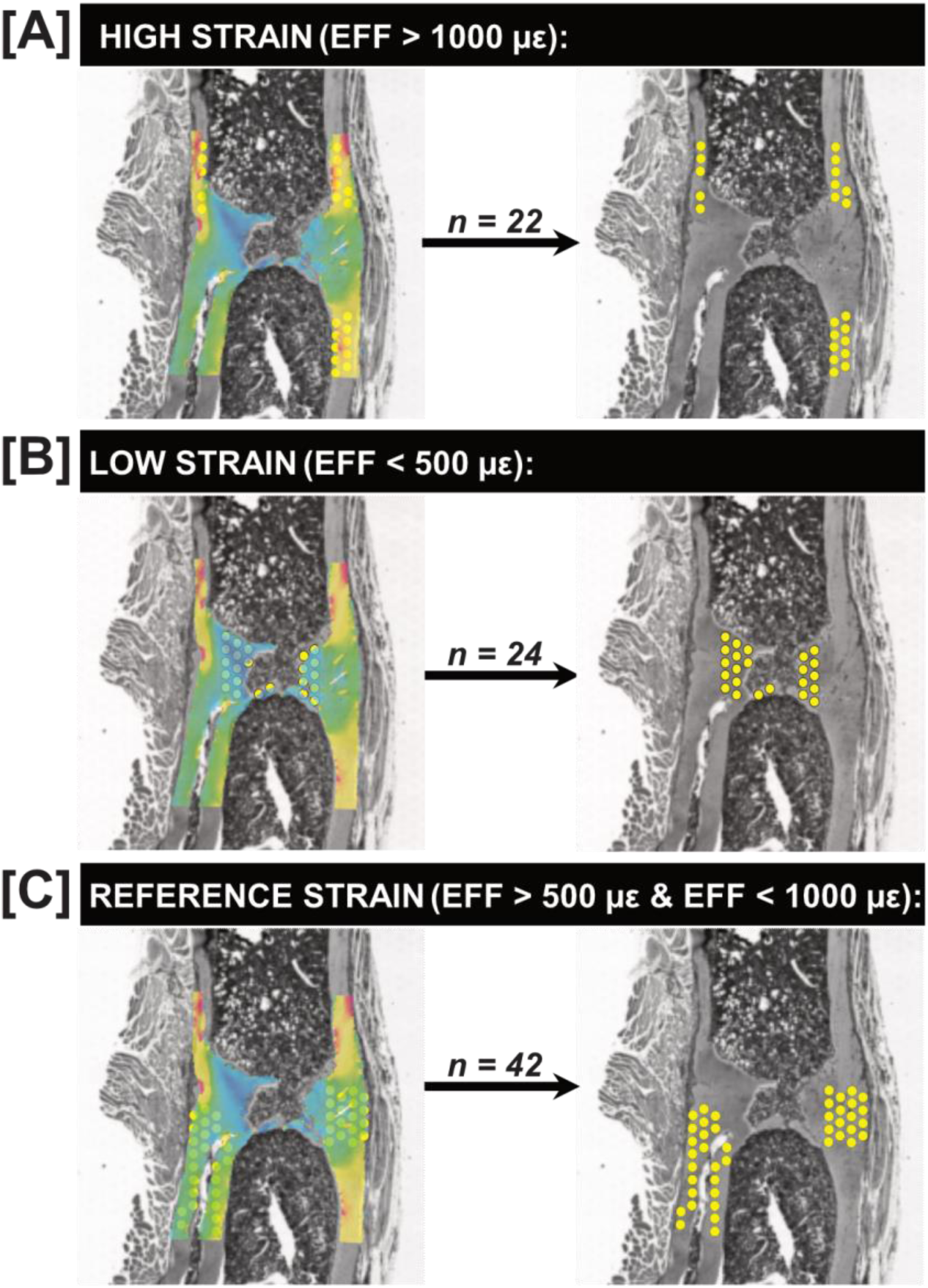
Classification of spots with respect to their local *in vivo* mechanical environment. Identification of transcriptomic responses at **[A]** sites of high strain (EFF > 1000 µε), **[B]** sites of low strain (EFF < 500 µε), and **[C]** sites corresponding to a reference strain region (EFF > 500 µε and EFF < 1000 µε). Element size in the 3D micro-FE model of the mechanical environment is 10.5 x 10.5 x 10.5 μm. Spot size of the spatial transcriptomics data is 55 μm. Effective strain (EFF) represents the mechanical environment.

**Fig. 8.**
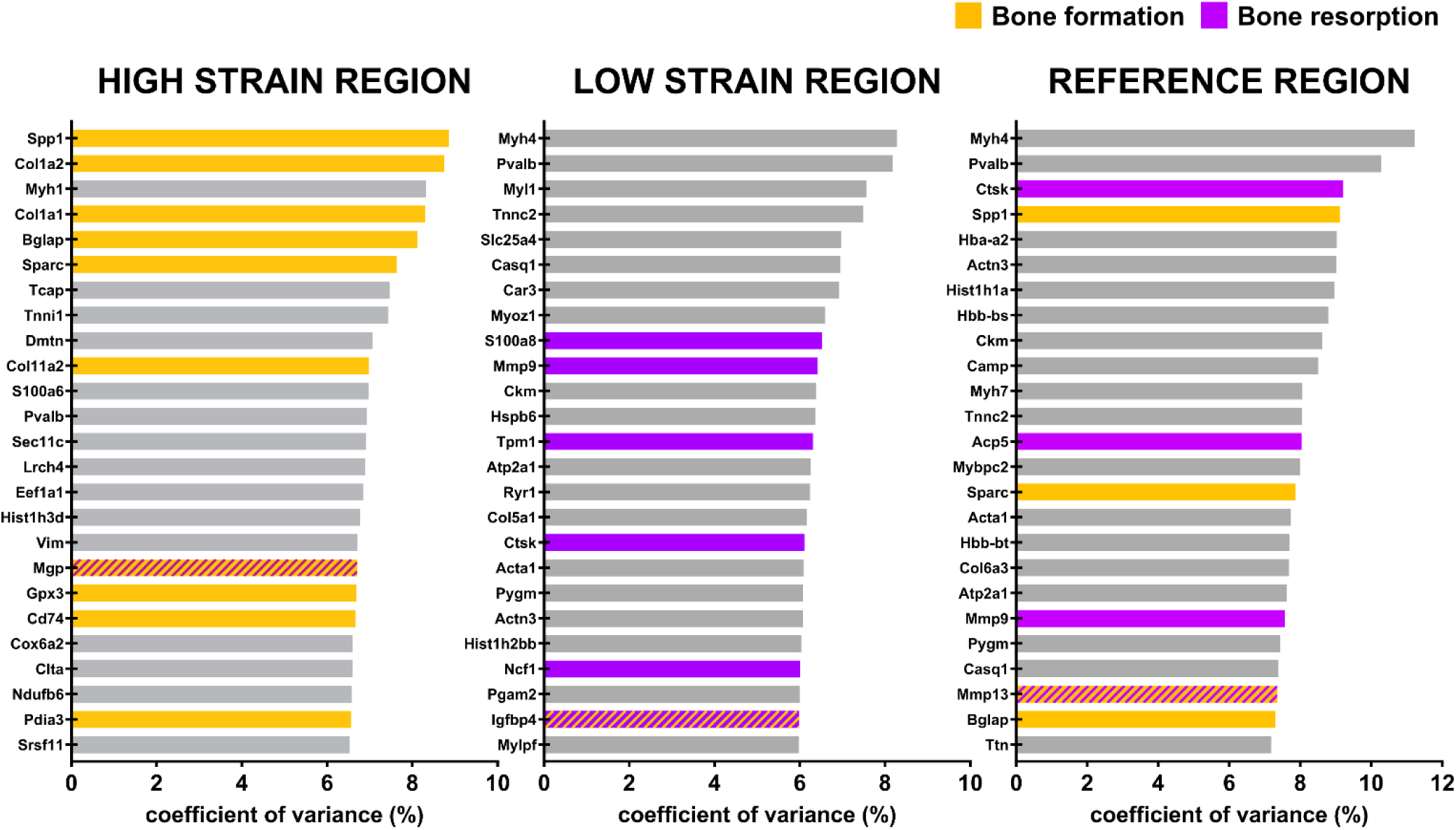
Use of the co-efficient of variance (CV) to analyze transcriptomic responses of cells with respect to their local *in vivo* mechanical environment. Data presented (n = 1 per group) corresponds to the mechanically loaded fracture site at 5 weeks post-surgery. Use of the co-efficient of variance (CV) to identify the top 25 genes of functional significance within each strain region. Genes associated with bone formation are identified in orange. Genes associated with bone resorption are identified in purple. Genes which are inhibitors of bone formation / resorption or genes which have roles in both bone formation / resorption are identified by alternating lines / hatches in orange and purple.

In comparisons of high strain vs. reference strain regions, 5146 genes were identified with sufficiently high expression to be included in a DEG analysis. Of these, 114 genes were differentially expressed (FDR-adjusted p value cutoff < 0.05; absolute log2 fold change > 0.5) with 108 genes upregulated and 6 genes downregulated (table S3). The following genes associated with bone formation were found to be upregulated: *Coq10a* (+2.2 fold) (*51*, *52*), *Myh2* (+2.0 fold) (*53*), *Sirt7* (+1.9 fold) (*54*), *Pdia3* (+1.0 fold) and *Col1a2* (+0.9 fold) (fig 9[A]; table S3). All remaining upregulated genes had no established roles in bone function or are ubiquitously expressed. Downregulated genes included the following gene associated with bone mineralization: *Sparc* (-0.8 fold); and the following genes associated with bone resorption: *Jdp2* (-1.6 fold) (*56*), *Mmp9* (-1.5 fold), *Acp5* (-1.4 fold) and *Ctsk* (-0.9 fold) (fig 9[A]; table S3). All remaining downregulated genes had no established roles in bone function. Gene-set enrichment analysis of differentially expressed genes identified the positive enrichment of signaling pathways associated with bone formation (FDR-adjusted p value < 0.05) (fig 9[B]; table S5).

**Fig. 9.**
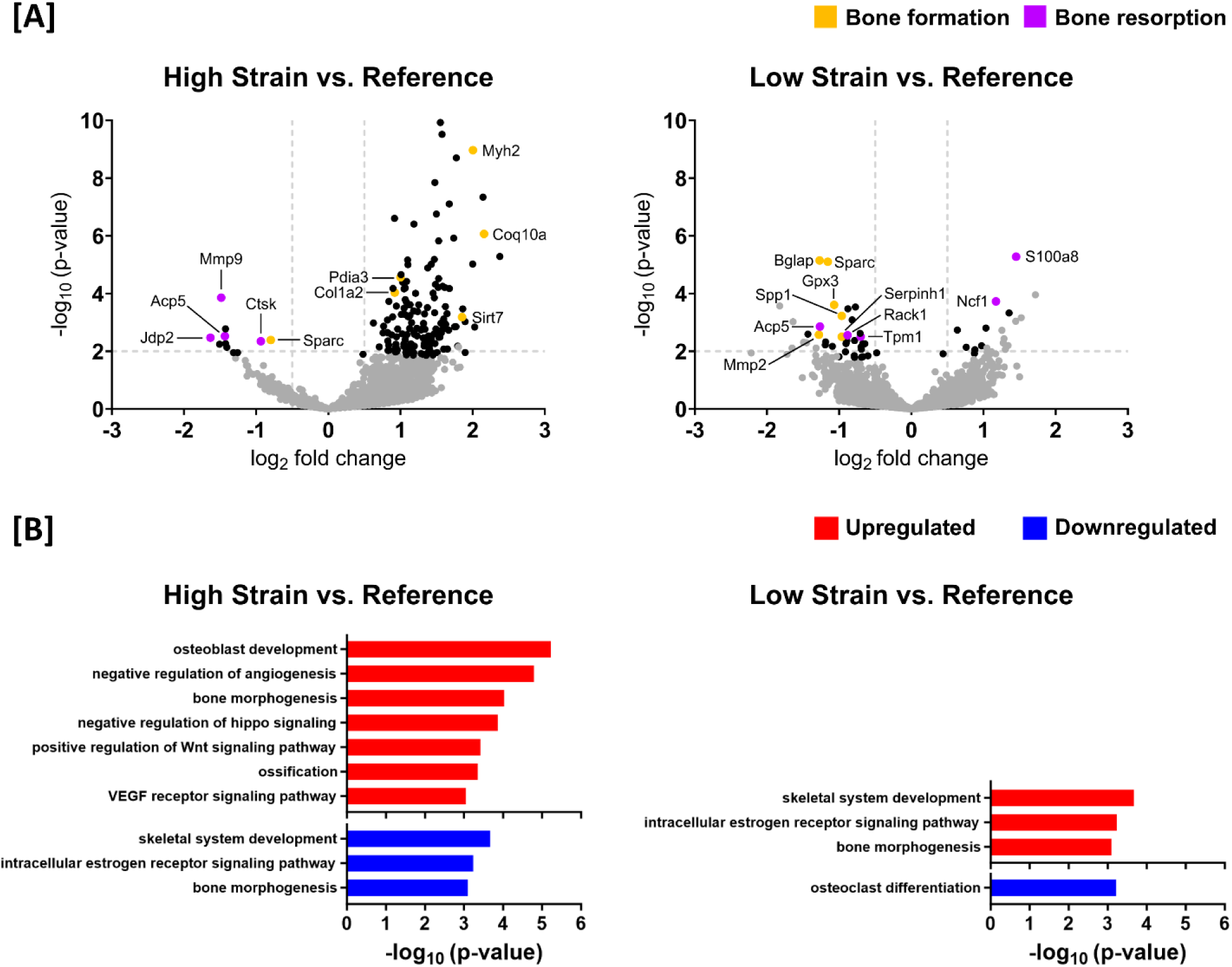
Use of differential gene expression analysis and gene-set enrichment analysis to analyze the transcriptomic responses of cells with respect to their local *in vivo* mechanical environment. Data presented (n = 1 per group) corresponds to the mechanically loaded fracture site at 5 weeks post-surgery. Genes associated with bone formation are identified in orange. Genes associated with bone resorption are identified in purple. Genes which are inhibitors of bone formation / resorption or genes which have roles in both bone formation / resorption are identified by alternating lines / hatches in orange and purple. **[A]**: Volcano plots to visualize differentially expressed genes (DEGs) between strain regions. (significance criteria: FDR-adjusted p value cutoff < 0.05 and an absolute log2 fold change > 0.5). Significant DEGs associated with bone formation are identified in orange and significant DEGs associated with bone resorption are identified in purple. Non-DEGs are represented in gray. **[B]:** Gene-set enrichment analysis performed in one-on-one comparisons between mean expression of high strain versus reference spots and low strain versus reference spots. The significant annotation terms were selected using an FDR-adjusted p value < 0.05. Only annotation terms relevant to fracture healing are presented.

In comparisons of low strain vs. reference strain regions, 5146 genes were identified with sufficiently high expression to be included in a DEG analysis. Of these, 19 genes were differentially expressed (FDR-adjusted p value cutoff < 0.05; absolute log2 fold change > 0.5) with 5 genes upregulated and 14 genes downregulated (table S4). The following genes associated with bone resorption were found to be upregulated: *S100a8* (+1.5 fold) and *Ncf1* (+1.2 fold) (fig 9[A]; table S4). All remaining upregulated genes had no established roles in bone function or are ubiquitously expressed. Downregulated genes included the following genes associated with bone formation: *Mmp2* (-1.3 fold), *Bglap* (-1.3 fold), *Sparc* (-1.2 fold), *Gpx3* (-1.1 fold) (*61*), *Spp1* (-1.0 fold) and *Serpinh1* (-1.0 fold) (*62*); and the following genes associated with bone resorption: *Acp5* (-1.3 fold), *Rack1* (-0.9 fold) (*66*) and *Tpm1* (-0.7 fold) (fig 9[A]; table S4). All remaining downregulated genes had no established roles in bone function.

## Discussion

The mechano-responsiveness of bone cells has been recognized for over 100 years, yet the underlying biological mechanisms remain not well understood. It has proven extremely difficult to investigate the role of local mechanical environments at fracture sites in driving site-specific cellular responses. In this study, we address this challenge by demonstrating the potential of our spatial transcriptomics based “mechanomics” platform in exploring the mechanoregulation of fracture healing. Using an established femur defect mouse model, we demonstrate that our cyclic mechanical loading protocol induces a strong anabolic response corroborated by time-lapsed micro-CT measurements and spatial transcriptomics profiling. Moreover, in associating gene expression profiles with their local mechanical environments, sites of high strain were associated with bone formation responses and sites of low strain were associated with bone resorption responses. Our spatial transcriptomics-based “mechanomics” platform thus presents unique opportunities to investigate fundamental questions within the field of bone mechanobiology: (i) which genes, cell populations and signaling pathways respond to local mechanical stimuli, (ii) how do the responses of these genes, cell populations and signaling pathways change as a function of the local mechanical environment, and (iii) how can the “mechanical dose” delivered by mechanical intervention therapies be optimized to achieve specific outcomes.

Micro-CT-based bone morphometric analysis confirmed the effect of cyclic mechanical loading with an enhanced osteogenic response producing much larger callus / bone volumes. Although limited to n = 2 mice per group, this is corroborated by our previous studies in female 20-week-old C57BL/6 mice (*11*), female 12-week-old and 35-week-old PolgA^D257A/D257A^ mice (*67*, *68*) and female 12-week-old BCR:PolgA ^D257A/D257A^ mice (*69*). Moreover, as illustrated in Figure 2, sites of bone formation (identified in orange) unequivocally predominate between weeks 3 – 5 in the mechanically loaded fracture sites.

Visualization of spatial gene expression at the fracture sites and differential gene expression analyses further underscored this osteogenic effect with upregulation of genes associated with matrix synthesis (*Col1a1*, *Col1a2* and the non-collagenous proteins: *Bglap*, *Sp7*) and mineralization (*Alpl*, *Dmp1*, *Phex*, *Car12*, *Smpd3*, *Phospho1*) (fig 5[B][C]; table S2). Chondrocyte markers were minimally present at both fracture sites as the timepoint analyzed post-fracture corresponded to the remodeling phase of the fracture healing process (fig S1). Despite the functional importance of members of the tumor necrosis family, significant differences in the expression of osteoprotegerin (OPG – encoded by *Tnfrsf11b*) and receptor activator of nuclear factor kappa-Β ligand (RANKL – encoded by *Tnfsf11*) were not present. Elevated expression of both OPG and RANKL has been reported in a tibial closed-fracture mouse model with levels gradually subsiding over a 4-week period relative to contralateral control tibias (*70*). However, in the present study, mechanical loading of the fracture site did not markedly alter the expression of OPG – an inhibitor of osteoclastogenesis. In agreement, markers of osteoclastogenesis (*Ncf1*, *Mmp9*) were found to be upregulated, and are likely the result of the coupling of the anabolic and catabolic phases of the response to mechanical loading.

The prevalence of osteocyte markers (*Phex*, *Dmp1*, *Mepe*, *Sost*) amongst the top differentially expressed genes is noteworthy and likely reflect the following processes at the fracture site: (i) mineralization driven by young osteocytes (*Phex*, *Dmp1*), (ii) regulation of mineralization by mature osteocytes (*Mepe*), and (iii) the progressive maturation of osteocytes encapsulated in the bone matrix (*Mepe*, *Sost*) (*71*). The upregulation of *Sost* – a potent inhibitor of Wnt signaling in osteoblasts – was unexpected. The expression of *Sost* is regulated by both biochemical (*72*, *73*) and mechanical (*38*) cues. In intact bones, *Sost* expression is downregulated with mechanical loading and upregulated with mechanical unloading (*38*). However, fewer studies have investigated the role of *Sost* in the context of fracture healing. Coates et al. conducted a comparison of the transcriptional profiles of intramembranous and endochondral ossification following fracture (*30*). In their model of intramembranous ossification in the mouse ulna, *Sost* was downregulated at 4 hours and 1 day following fracture relative to the intact contralateral ulna. In their model of endochondral ossification in the mouse femur, *Sost* was downregulated at 4 hours, 1 day, 3 days, 7 days and 14 days following fracture relative to the intact contralateral femur. The bone formation mechanism (that is, intramembranous vs. endochondral ossification) within each of these models is determined by the mechanical stability of the injury site. This demonstrates that the regulation of *Sost* at fracture sites is not exclusively mechanical in nature. Intriguingly, the presence of an alternative mechanism of Wnt regulation is suggested in our results by the upregulation of *Wnt7b* and downregulation of *Sfrp4*. Expression of *Wnt7b* is reported to be minimal in adult bone – but induced in response to mechanical loading (*74*) or following fracture (*30*). Downregulation of the Wnt antagonist *Sfrp4* is similarly reported to be mechanically-induced in rodent models of bone adaptation (*41*, *42*). Neither gene has been well-studied in the context of mechanically-driven fracture healing and thus warrants further investigation as a potential mechano-sensitive regulatory mechanism of Wnt signaling. Nevertheless, it should be noted that our spatial transcriptomics data is limited to n = 1 per group – thus our findings may not be representative of larger data sets. In the same way that each mouse in our study receives an individualized or “patient-specific” loading intervention, the fracture healing response and the response to mechanical loading are variable processes that differ from subject-to-subject.

In associating molecular pathways at the cellular scale with their local mechanical *in vivo* environment within a single histological section, cells in regions of high strain were found to express genes involved in bone formation responses: upregulation of *Coq10a*, *Myh2*, *Sirt7*, *Pdia3* and *Col1a2*, downregulation of *Jdp2*, *Mmp9*, *Acp5* and *Ctsk*, and positive enrichment of “*osteoblast development*”, “*ossification*”, “*positive regulation of Wnt signaling pathway*” and “*negative regulation of hippo signaling*”; whereas cells in regions of low strain were found to express genes involved in bone resorptive responses: upregulation of *S100a8* and *Ncf1* and downregulation of *Mmp2*, *Bglap*, *Sparc*, *Gpx3*, *Spp1* and *Serpinh1*. This finding is in agreement with the fundamental principle of Wolff’s Law on the capacity of bone to functionally adapt to its mechanical environment. However, we are yet to establish the mechanobiological mechanisms governing these strain-specific cellular responses as this would necessitate a more comprehensive dataset. Indeed, it cannot be ruled out that the cellular responses observed are (to a partial extent) site-specific responses or biochemically-driven responses independent of the local mechanical environment. Nevertheless, this demonstration within a single histological section of bone underscores the potential of our platform to develop a molecular based understanding of the mechanoregulation of fracture healing. Indeed, the ability to analyze spatially-resolved omics data with respect to the local *in vivo* mechanical environment is a notable achievement within the field of bone mechanobiology.

In assessing the merits of our platform, original features, opportunities for further optimization and limitations were considered. There are three original aspects to our work: (i) no comparable platform exists in the field which permits the spatial integration of CT bone morphology data, 3D mechanical environments and gene expression data from a single fracture site, (ii) the generation of spatially resolved transcriptomics data to investigate bone mechanobiology, and (iii) the analysis of gene expression as a function of the local mechanical environment. The demonstration of the latter analysis is the most impactful feature of our work. Previous studies have largely assumed homogeneous strain environments at skeletal sites as the technology has simply not existed to analyze gene expression as function of local mechanical environments (*75*). It thus represents a major technological achievement as it surpasses the current state-of-the-art in the field of bone mechanobiology. Furthermore, our platform has broad applications within the field of bone mechanobiology. In mouse models, the three most common skeletal sites at which mechanobiological studies have been conducted are the femur, the vertebra (*76*) and the tibia (*77*). These studies have investigated the mechanobiology of either bone adaptation or fracture healing. By replacing the femur defect mouse model in our platform with one of these other established mouse models in the field, our platform can be applied to investigate the mechanobiology of bone at these sites. All other components of our platform (micro-CT imaging, mechanical loading, micro-FE modelling, spatial transcriptomics) can be applied at any of these skeletal sites. Indeed, the platform is also adaptable for use with tissue engineered bone constructs.

The spatially resolved characterization of the transcriptome of a fracture site presented herein is a notable feature. Historically, experimental methodologies within the field have presented a trade-off between the use of omics technologies to sequence the genome, transcriptome or proteome from dissociated specimens vs. the use of immunohistochemistry or *in situ* hybridization to localize a predetermined subset of molecules within intact tissues. With the advent of spatial molecular profiling technologies, spatially contextualized maps of the diverse landscape of cell types and their functions can be captured. However, the use of spatial transcriptomics in bone presents challenges due to the mineralized nature of bone. The need to maintain a delicate balance between tissue decalcification and RNA preservation has constrained the use of spatial transcriptomics in bone studies. However, we have established and made available a protocol for the use of the Visium Spatial Gene Expression assay with FFPE bone sections (*13*). The success of our protocol is evident in quality control measures. In measures of both unique transcripts and unique genes, the median numbers for our data were notably higher than those reported for bone sections in the literature (*78*). In terms of reproducibility, each tissue- and strain-specific spot within a spatial transcriptomics section can be considered a technical replicate of the local *in vivo* environment. All spots within a section are subjected to the same sample preparation and processing conditions, reducing technical variation introduced by differences in the processing of different samples. Moreover, comparing spots from different regions within the same tissue section can provide a means of internal validation and control. For example, in our analysis of strain regions within the mineralized tissue, regions of high / low strain are compared against a reference strain region within the same section. Although beyond the scope of this publication, our experiments with the Visium Spatial Gene Expression assay also permitted preservation of muscle and marrow at the fracture site. The technique thus provides opportunities to investigate the crosstalk between different cell populations and its roles in driving the healing response at the fracture site. Notably, genes without well-defined roles in fracture healing also featured prominently in our differential gene expression analyses underscoring the transformative potential of spatial profiling technologies in unravelling molecular pathways and mechanisms.

The platform does present opportunities for further optimization and is not without limitations. To associate transcriptomic responses of cells to their local *in vivo* mechanical environment, a 2D histological section was visually aligned within the 3D micro-CT derived mechanical environment. However, the use of an approach based on visual assessment is both labor intensive and error prone. Machine learning driven 2D-to-3D registration techniques can potentially be incorporated into the platform to perform this task. Moreover, the transcriptomics data generated using the Visium Spatial Gene Expression assay is not at single-cell resolution. Instead, the capture area on each Visium slide consists of a grid of 55 µm diameter spots with the center of each spot positioned approximately 100 µm from the center of adjacent spots. Each spot may thus overlap the boundaries between tissue structures or strain regions and may necessitate the exclusion of specific spots. Calluses with trabecular structures pose a specific challenge as the grid pattern of the Visium capture areas may not be optimally positioned to capture the transcriptomic response within each trabecular strut. Recent advancements in omics technologies have led to the availability of techniques for generating spatially-resolved omics data at single-cell resolution (such as the Visium HD slide from 10x Genomics). Integration of such techniques into our platform represents the next logical progression in its evolution. In addition, the RNA quality of transcriptomics data generated from formalin-fixed paraffin embedded samples is generally inferior to fresh frozen samples. Protocols for the use of fresh frozen bone samples with Visium Spatial Gene Expression assay are yet to be established and future applications using our platform should consider the use of fresh frozen samples. Finally, the micro-FE models of the mechanical environment are based on supra-physiological loading applied to the fracture site and do not consider the physiological loading applied during functional activities. Direct measurements of the mechanical environment via implanted sensors could provide quantification of the physiological loading at the fracture site (*79*) – but implementation of such sensors in mouse models has proven challenging.

In conclusion, we present an experimental platform to perform spatially-resolved analysis of the transcriptomic responses of cells with respect to their local *in vivo* mechanical environment. Given the limited understanding of the cellular and molecular mechanisms governing the mechanobiology of bone repair, the platform – especially when adapted to function at single-cell resolution – has the potential to address the fundamental open question within the field: Which cell populations and signaling pathways sense and respond to local mechanical stimuli? Insights into the mechanoregulation of fracture healing will have implications for the broader translation of mechano-therapeutics to clinical settings, with the potential to identify strategies and discover mechano-responsive targets to enhance repair in compromised healing environments.

## Materials and Methods

### Experimental Design

Our objective is to demonstrate the potential of our spatial-transcriptomics based mechanomics platform in investigating the mechanobiology of fracture healing. As illustrated in Figure 1, the platform consists of: (i) an established femur defect mouse model (*11*), (ii) established *in vivo* micro-CT imaging protocols and analyses (*9*), (iii) an established osteogenic cyclic mechanical loading approach (*11*, *12*), (iv) an established spatial transcriptomics approach for bone tissue (*13*) and (v) an established *in silico* micro-FE modelling approach (*12*).

### Ethics statement

All mouse experiments were performed in accordance with relevant national regulations (Swiss Animal Welfare Act, TSchG, and Swiss Animal Welfare Ordinance, TSchV)) and authorized by the Zürich Cantonal Veterinary Office (approved license number: ZH229/2019; Kantonales Veterinäramt Zürich, Zurich, Switzerland).

### Mouse line

Our lab has previously established a bone cell reporter (BCR) mouse model using CRISPR/Cas9 technology to label osteoblast (Integrin binding sialoprotein - Ibsp) and osteoclast-specific targets (tartrate-resistant acid phosphatase (TRAP) type 5 - Acp-5) with fluorescent proteins (eGFP, mCherry) (*80*). All mice were bred, monitored and maintained under specific pathogen free (SPF) conditions at the ETH Phenomics Centre, ETH Zürich (12 h:12 h light-dark cycle, ad libitum access to maintenance feed and water).

### Femur defect model

Female 12-week-old BCR mice (n = 4) received mid-diaphyseal femoral defects (0.68 ± 0.04 mm) using an established osteotomy surgical protocol (*11*). Briefly, an external fixator (Mouse ExFix, RISystem, Davos, Switzerland) was positioned at the craniolateral aspect of the right femur using four mounting pins and the defect was created using a 0.66 mm Gigli wire saw. Pre-operative analgesia (25 mg/L, Tramal®, Gruenenthal GmbH, Aachen, Germany) was provided via the drinking water two days before surgery until the third post-operative day. Anesthesia for all animal procedures (surgery, *in vivo* imaging, mechanical loading) was achieved using isoflurane (induction/maintenance: 5%, 2–3% isoflurane/oxygen).

### *In vivo* micro-CT imaging

*In vivo* micro-CT imaging of the fracture site between the two inner screws of the external fixator was performed weekly in all mice (weeks 0-5; vivaCT 80, Scanco Medical AG, Brüttisellen, Switzerland) (10.5 μm resolution, 55 kVp, 145 µA, 350 ms integration time, 500 projections per 180°, 21 mm field of view (FOV), scan duration ca. 15 min). To avoid motion artifacts during scanning, a custom-designed holder was used to secure the external fixator (*11*).

Registration of time-lapsed *in vivo* images permits visualization of sites of bone formation, quiescence and resorption (*76*). Reconstructed micro-CT images of each mouse were registered sequentially using an established algorithm (*9*); proximal and distal cortices were registered separately at time-points where bridging of the fracture site was not present as minor relative displacements between proximal and distal cortices were observed to occur. Images were Gaussian filtered (sigma 1.2, support 1) and bone volumes (bone volume—BV) were computed (threshold: 395 mg HA/cm^3^) in four non-overlapping volumes of interest (VOIs): the defect center (DC), the defect periphery (DP), the existing fracture cortices together with the medullary cavity (FC) and the cortex periphery (FP) (fig 3) (*9*). Bone morphometric indices (bone volume/total volume— BV/TV, bone formation rate – BFR, bone resorption rate – BRR) were evaluated within each VOI. Bone volumes were normalized with respect to the central VOIs (DC, FC) which represent the total volume (TV) of intact bone (*9*): thus, DC/DC, DP/DC, FC/FC, FP/FC. Defect sizes (h) were calculated using the following formula: h = 2DC / (CSA_P + CSA_D) where DC is the defect volume at week 0 and CSA_P and CSA_D represent the proximal and distal cross-sectional areas respectively, which are situated directly adjacent to the fracture site.

### Mechanical loading

Following bridging of the fracture site at 3 weeks post-surgery, the mice received either individualized cyclic mechanical loading (*11*, *12*) via the external fixator (n = 2, 8-16N, 10Hz, 3000 cycles) or 0N sham-loading (n = 2). Loading is applied 3 times per week. Detailed descriptions of the protocols can be found in the literature (*11*, *12*). In brief, the mechanical loading applied at each time point is based on computation of the strain distribution within the fracture site of each mouse and the scaling of the strain distribution to achieve a pre-defined median target strain. This is implemented in real-time within a single anesthetic session each week as follows: directly following *in vivo* micro-CT imaging, the CT data is reconstructed and a high-resolution *in silico* finite element (FE) model of the fracture site generated. The FE model simulates axial compression (48-core Intel(R) Xeon(R) Platinum 8168 CPU @ 2.70GHz) and a histogram of the strain distribution is generated. The strain distribution is subsequently scaled by the applied load to achieve a pre-defined median target strain. Furthermore, to assess if the loading poses a structural failure risk at the fracture site, the simulation identifies voxels which exceed more than 10000 µε. If more than 100 voxels exceed this threshold, the loading value is reduced by 1N. This optimized load is then applied to the mouse. As the optimized load is based on the *in vivo* micro-CT image and *in silico* simulation at each weekly time-point, loading is individualized to each mouse such that the induced median strain in all loaded mice is of comparable magnitude (*12*).

### Spatial transcriptomics

All mice were euthanized at 10 hours following the final cyclic mechanical loading session at 5 weeks post-surgery. Spatially resolved transcriptomics analyses were performed on explanted femurs (n = 1 per group) using the Visium Spatial Gene Expression for formalin-fixed paraffin-immediately fixed in 10% neutrally buffered formalin for ca. 16 – 24 hours at 4°C, decalcified in 12.5% EDTA (pH 7.5) for 10 days at 4°C, placed in a tissue processor and embedded in paraffin. Protocol details have been published separately using the section from the Control mouse (*13*). To assess the RNA quality, RNA was extracted from each sample (Qiagen RNeasy FFPE Kit, Hilden, Germany) and the DV 200 value (that is, the percentage of total RNA fragments >200 nucleotides) of each sample was evaluated (Agilent 4200 TapeStation, Waldbronn, Germany). As per the manufacturers’ instructions, only samples with a DV 200 > 50% were selected for spatial gene expression analyses. 5 µm longitudinal sections from each explanted femur (n = 1 Control, n = 1 Loaded) were placed onto 6.5 x 6.5 mm capture areas on a Visium Spatial Gene Expression slide. Sections were subsequently deparaffinized, subjected to hematoxylin and eosin (H&E) staining, imaged and decrosslinked in accordance with manufacturers’ instructions (10x Genomics, CG000409, Rev D). Sections were probe hybridized with 20551 genes targeted (10x Genomics, Visium Mouse Transcriptome Probe Set v1.0) and spatial transcriptomics libraries were prepared. Libraries were sequenced on an Illumina NovaSeq6000 System (Illumina, San Diego, United States) at a sequencing depth of approximately 75 – 120 million reads per sample.

### Analysis of sequencing data

Demultiplexing and manual alignment of the sequencing data to the histological image were performed using the SpaceRanger analysis pipeline (10x Genomics, version 2.0.0) and Loupe Browser (10x Genomics, version 6.2.0) respectively. Further downstream data analyses and visualization were performed in R (version 4.3.1) using Seurat (version 4.4.0). Spots were excluded by filtering each sample separately based on the number of UMIs (nCount_Spatial >= 500) and the number of genes (nFeature_Spatial >= 250).

#### Regions of interest

Regions of interest (ROI) were defined in Loupe Browser by selecting spots which corresponded to specific structures in the H&E stained histological images. Barcodes corresponding to all spots in each region of interest are provided in Supplementary Table S1. In comparisons between Control and Loaded sections, ROIs were defined at the fracture site encompassing all bone spots between the two middle pins (fig 5; Control: n = 131 spots, Loaded: n = 146 spots). The entire region between the two middle pins was selected as in the loaded mouse this region is subjected to loading. Furthermore, in the Loaded section, ROIs were defined based on the local mechanical environment: high strain region (> 1000 µε), low strain region (< 500 µε) and reference strain region (> 500 µε and < 1000 µε).

#### Differential gene expression analysis

Differential gene expression analyses were performed using DESeq2 by implementing scaling normalization through deconvolving size factors using scran. In addition, null hypothesis testing was conducted using the likelihood ratio test (*81*). Lists of differentially regulated genes for the different comparisons are provided in Supplementary Tables S2–S4. Differentially expressed genes were selected using an FDR-adjusted p value cutoff < 0.05 and an absolute log2 fold change > 0.5. To compare gene expression profiles at sites of high and low strain, the co-efficient of variance (CV) was used to identify genes of functional significance across all spots within a region.

#### Gene enrichment analysis

Gene-set enrichment analysis was performed using Generally Applicable Gene-set Enrichment (GAGE; Bioconductor version 3.18). For functional annotation, gene sets from org.Mm.eg.db, a genome-wide annotation package for mouse, were utilized. The analysis was performed via one-on-one comparisons of mean gene expression between high strain vs. reference regions and low strain vs. reference regions. The significant annotation terms were selected using an FDR-adjusted p value < 0.05. Lists of enriched pathways are provided in Supplementary Tables S5 and S6.

### *In silico* micro-FE modelling

Micro-FE analyses based upon the registered *in vivo* micro-CT images were used to simulate axial compression and generate tissue-scale 3D maps of the mechanical environment (*12*, *82*). In brief, converted to linear hexahedral elements to generate an FE mesh. Grayscale values of the voxels were then converted from density (mg HA/cm^3^) to Young’s moduli (GPa) (*83*). Regions of soft tissue were assigned a Young’s modulus of 0.003 GPa (*84*). In addition, the marrow cavity of the femur was capped on the top and bottom slices of the image stack with a plate of 20 GPa to prevent edge effects due to the presence of soft tissue. To map the mechanical environment at the fracture site, uniaxial loading was simulated by applying a 1% compressive displacement to the top slice in the axial direction whilst the bottom slice was held fixed. ParOSol - a linear micro-FE solver - was used to solve each finite element simulation and compute the mechanical environment (*85*). Effective strain (EFF), which combines both volumetric and deviatoric strains (and drives fluid movement and direct strain, respectively), represented the mechanical environment (*9*, *82*). The results of the simulation were scaled as follows:

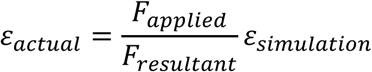

where *ε*_*simulation*_ is the effective strain result based on the simulation of uniaxial loading, *F*_*resultant*_ is the sum of the reaction forces of all the nodes of the uppermost surface, *F*_*applied*_ is the cyclic loading force applied and *ε*_*actual*_ is the strain induced under the applied force (*82*).

### Mechanomics

Histological sections on which spatial transcriptomics were performed were visually aligned within the 3D maps of the mechanical environment to correlate spatially resolved gene expression profiles with their local in vivo mechanical environments (fig 6; ParaView version 5.7.0). Maps of the mechanical environment were sub-divided into high strain (EFF > 1000 µε) and low strain (EFF < 500 µε) regions, and the corresponding gene expression profiles were analyzed with respect to a reference strain region (EFF > 500 µε and EFF < 1000 µε).

## Acknowledgments

The authors gratefully acknowledge the ETH Phenomics Centre (EPIC) of ETH Zürich, and particularly Susanne Freedrich, for assistance with the *in vivo* experiments.

Spatial transcriptomics was performed at the Functional Genomics Center Zürich (FGCZ) of the University of Zürich and ETH Zürich.

Tissue processing and paraffin embedding were performed at ScopeM at ETH Zürich.

Sectioning, staining and imaging were performed at ScopeM at ETH Zürich and at the Institute of Pathology and Molecular Pathology at the Universitäts Spital Zürich (USZ).

## Funding

European Research Council - Horizon 2020 grant ERC-2016-ADG-741883

European Union Marie Skłodowska-Curie Actions - Horizon 2020 grant 101029062

MechanoHealing-MSCA-IF-2020

Swiss National Science Foundation (IZCOZ0_198152/1, COST Action GEMSTONE)

## Author contributions

Conceptualization: NM, EW, RM

Methodology: NM, DG, EW, RM

Investigation: NM, GK, EW, RM

Analysis: NM, AS, FCM

Data Curation: NM, AS

Funding Acquisition: NM, EW, RM

Supervision: EW, RM

Visualization: NM, AS

Writing—original draft: NM

Writing—review & editing: All.

## Data and materials availability

All data needed to evaluate the conclusions in the paper are present in the paper and/or the Supplementary Materials. Original spatial transcriptomics data generated for this article has been deposited in the Gene Expression Omnibus (GEO) database (accession code: XXXXXXXX).

## Supplementary Materials

**Supplementary Fig. S1.**
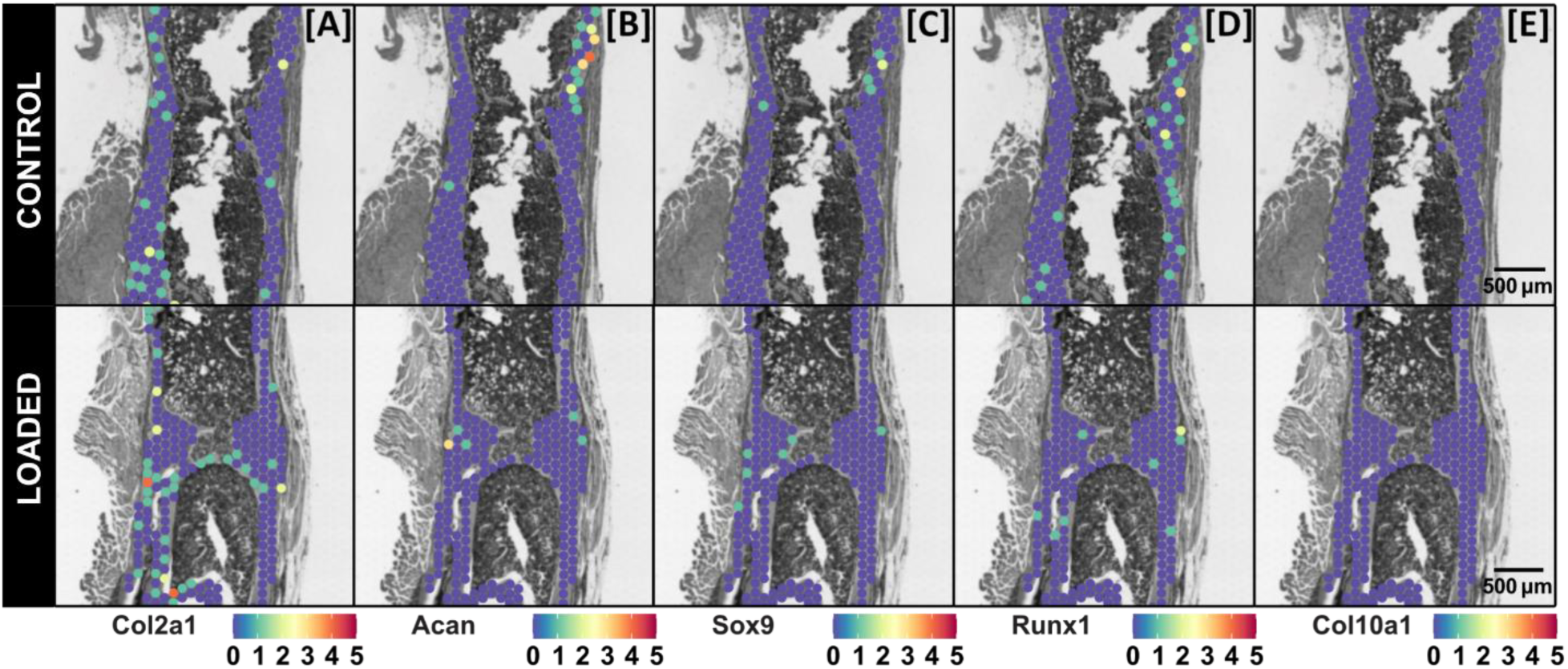
Spatial gene expression maps of selected chondrocyte markers at the fracture site of Control and Loaded mice. Visualization of the spatial expression patterns of chondrocyte markers: **[A]** *Col2a1*, **[B]** *Acan*, **[C]** *Sox9*, **[D]** *Runx1* and **[E]** *Col10a1* within the fracture sites of Control and Loaded mice are presented. Each legend denotes the normalized expression of each gene. Data presented (n = 1 per group) corresponds to samples at 5 weeks post-surgery. 3D visualizations of the morphology of these Control and Loaded fracture sites are presented in Figure 2. Spatial transcriptomics spot size = 55 μm.

**Supplementary Fig. S2.**
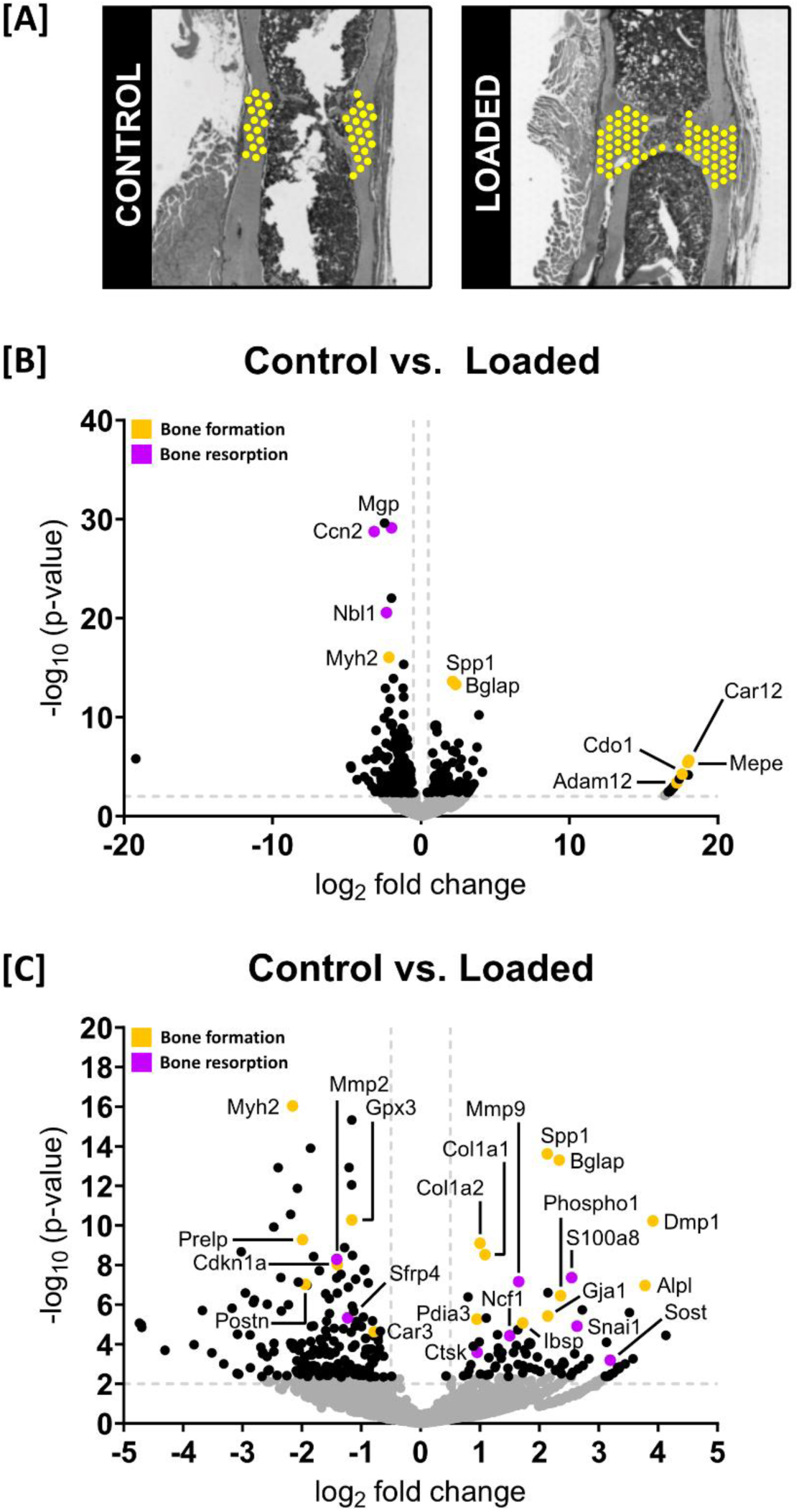
Gene expression profiling in DC + DP regions of Control vs. Loaded fracture sites. Data presented (n = 1 per group) corresponds to samples at 5 weeks post-surgery. **[A]**: Areas of interest defined in each fracture site for the analysis. **[B]**: Volcano plot to visualize differentially expressed genes (DEGs) (significance criteria: FDR-adjusted p value cutoff < 0.05 and an absolute log2 fold change > 0.5). Significant DEGs associated with bone formation are identified in orange and significant DEGs associated with bone resorption (or are inhibitors / antagonists of bone formation) are identified in purple. Non-DEGs are represented in gray. **[C]**: Magnified view of volcano plot to highlight DEGs of interest.

**Supplementary Table S1.** Barcodes corresponding to all spots of the Control and Loaded bone fracture sites.

**Supplementary Table S2.** Differential gene expression analyses (DEG) between Control and Loaded fracture sites.

**Supplementary Table S3.** Differential gene expression (DEG) analysis of high strain region with respect to reference region.

**Supplementary Table S4.** Differential gene expression (DEG) analysis of low strain region with respect to reference region.

**Supplementary Table S5.** Gene-set enrichment analysis between mean expression of high strain versus reference strain regions.

**Supplementary Table S6.** Gene-set enrichment analysis between mean expression of low strain versus reference strain regions.

